# Improving Target–Adverse Event Association Prediction via Inverse Node Frequency Transformation to Mitigate Knowledge Graph Topological Bias

**DOI:** 10.64898/2026.11.19.767791

**Authors:** Terence M. Egbelo, J. Charles G. Jeynes, Zeyneb Kurt, Rahul V. Veettil, Valerie J. Gillet

## Abstract

**Motivation:** Identifying significant associations between therapeutic protein targets and adverse events (AEs) is a critical component of drug safety assessment. To accelerate the process, machine learning methods have been developed to predict these associations as edges in network data integrating molecular interactions and other key biomedical knowledge. However, these networks often suffer from topological imbalances where a minority of “hub” nodes determine the predictive signal. This “topological” or “degree bias” can lead to poor performance on novel or understudied targets where safety inference is most needed.

**Results:** We introduce Inverse Node Frequency (INF), a generally applicable node degree-based approach for mitigating topological bias. Employing an enriched biomedical knowledge graph and our own carefully curated target-adverse event associations, we apply INF as a normalisation procedure on a class of graph-derived features called metapaths and evaluate their resulting target-AE association prediction performance. Our results demonstrate that INF significantly improves the true positive rate (TPR/recall) for low-degree targets compared to the untransformed metapath features as well as a popular established normalisation method, while also achieving highly competitive hits@*k*, a ranking-based metric for knowledge graph-based inference. We also evaluated INF on an external set of target-AE associations never before used for modelling, on which INF also showed superior resilience compared to the two benchmarks.

**Availability:** https://github.com/TERENTIVS/INF_KGC; archived at Software Heritage, swh:1:rev:09ff3525acb878c37a58d744b32ff11e4 5b14fad

**Supplementary information:** Supplementary data are provided with this submission.

## 1. Introduction

Adverse events (AEs), also known as adverse drug reactions (ADRs), are unintended and harmful reactions resulting from pharmaceutical intervention. These reactions represent a significant public health burden and a leading cause of drug withdrawals from the market (Boland et al 2016).

Drug-induced toxicities may be categorised into chemical-based (e.g. cellular damage or genetic toxicity via reactive metabolites) and target-based effects (Roberts 2018). Target-based safety liabilities may be further divided into on-target and off-target effects (Bowes et al 2012). On-target toxicity occurs when the candidate interacts with the correct target but at an inappropriate intensity or in a non-target tissue. Off-target toxicity arises when a drug interacts with unintended proteins, for example, the hERG potassium channel, blockage of which is associated with life-threatening cardiac arrhythmias (Sanguinetti et al 1995).

Consequently, the pharmaceutical industry has increasingly adopted a “target-centric” perspective on safety. Target Safety Assessment (TSA) has emerged as a cornerstone of preclinical derisking, where the safety profile of a putative therapeutic target is evaluated prior to the development of lead compounds (Blomme and Will 2016, Roberts 2018). Historically, this has relied on in vitro profiling against panels of “safety targets”. However, these panels are limited in scope and cannot feasibly cover the thousands of proteins in the human proteome. To scale these insights, computational methods have been developed to systematically link proteins to AEs. For example, the methodology by Kuhn et al (2013) exploits statistical associations between known drug-target and drug-AE associations to establish putative target-AE relationships (notably implemented at scale by the T-ARDIS database; Galletti et al 2021).

Nevertheless, statistics-based approaches rely on the availability of drug-AE association data (e.g. from drug packaging or reporting systems like FAERS) and knowledge of drug-target interactions in order to assert target-AE associations. For the vast majority of potential targets – for which no approved drugs yet exist – this “bridge” is unavailable, rendering statistics-based approaches inapplicable.

Recent advances in systems biology seek to fill this gap using network-based machine learning (ML). The few notable works in this direction so far, such as by Galletti et al (2022) and Park et al (2023) leverage techniques including network diffusion and graph representation learning on protein-protein interaction (PPI) networks and added-on drug-target relationships to train AE association classifiers for as-yet undrugged targets.

Despite the promise of network-based approaches, a critical challenge remains largely unaddressed, namely topological imbalance, or “degree bias”. This is the tendency of a small minority of “hub” nodes to accumulate a disproportionately high number of edges. With associations involving understudied or high-specificity entities forming the minority, the imbalance can lead to misleading performance metrics and poor generalisation (Bonner et al 2022a; Zietz et al 2024). In the context of target safety, hubs would comprise well-researched proteins with extensive known safety profiles; out-of-distribution failures would then occur precisely on those low-data proteins for which safety inference is most acutely needed.

In this study, we leverage a rich biomedical knowledge graph (KG) tailored to target-AE association prediction and propose a topological bias mitigation concept we term Inverse Node Frequency (INF). We apply it as a transformation of the path-based features (“metapaths”) of the interpretable KG completion approach introduced by Sun et al (2011). This method’s demonstrated yet overlooked promise in drug discovery is evident in the limited set of prior works which employ it, most notably Fu et al (2016) and Himmelstein et al (2015, 2017).

We demonstrate that the INF transformation maintains both a high True Positive Rate (TPR; recall) and Hits@*k* (a ranking-based KG completion metric) for low-degree targets where established transformations struggle. We also confirm these findings on an externally sourced target-AE association set compiled by Soldatos et al (2018) which has never before been used in this prediction task.

## 2. Materials & Methods

### 2.1 Collection of knowledge graph source data

The knowledge graph utilised in this study inherits most of its structure and content from the Drug Repurposing Knowledge Graph (DRKG) originally reported by Ioannidis et al (2020). The DRKG integrates information from a variety of biomedical resources.

The largest components of the DRKG are Hetionet (Himmelstein et al 2017), STRING (Szklarczyk et al 2016), DrugBank (Wishart et al 2018) and the Global Network of Biomedical Relationships (GNBR; Percha and Altman 2018).

For the purposes of target safety assessment (TSA), Evotec made specific modifications to update the content of the original DRKG.

- To maximise the graph’s relevance to human biology, information from animal studies was removed. This was achieved by removing all paths originating from non-human Gene nodes.
- Hetionet triples representing Gene Ontology (GO) annotations were replaced and updated with data from the most recent GO release (Gene Ontology Consortium 2019).
- Protein targets In the graph, are represented by their coding genes’ identifiers in the Entrez Gene database. Consequently, the node type for protein targets is labelled “Gene”.
- Relation (edge) types are labelled by concatenating the head entity type and the tail entity type (e.g., “Compound_Gene”). This design decision prevents the distinction of multiple association types between the same entity types.

### 2.2 Curation of the target-adverse event associations

Training the KG completion classifier required curating a set of ground truth target-AE associations and integrating them into the KG. These pairings represent causal relationships between target modulation and adverse event manifestation. These associations were compiled at Evotec through two complementary routes designed to minimise false positives (see Figure 1). Knowledge-guided associations were drawn from (1) the Open Targets platform (Ochoa et al 2023) and (2) in-house using custom NLP tools. Computationally derived associations were sourced from the T-ARDIS database (Galletti et al 2021), which applies Kuhn et al’s framework to large-scale AE case-report data (FAERS, MEDEFFECT) and curated data (SIDER and similar; Kuhn et al 2010) (Figure 1). The final ground truth set comprised 6,657 associations spanning 473 unique targets and 537 AEs.

**Figure 1:**
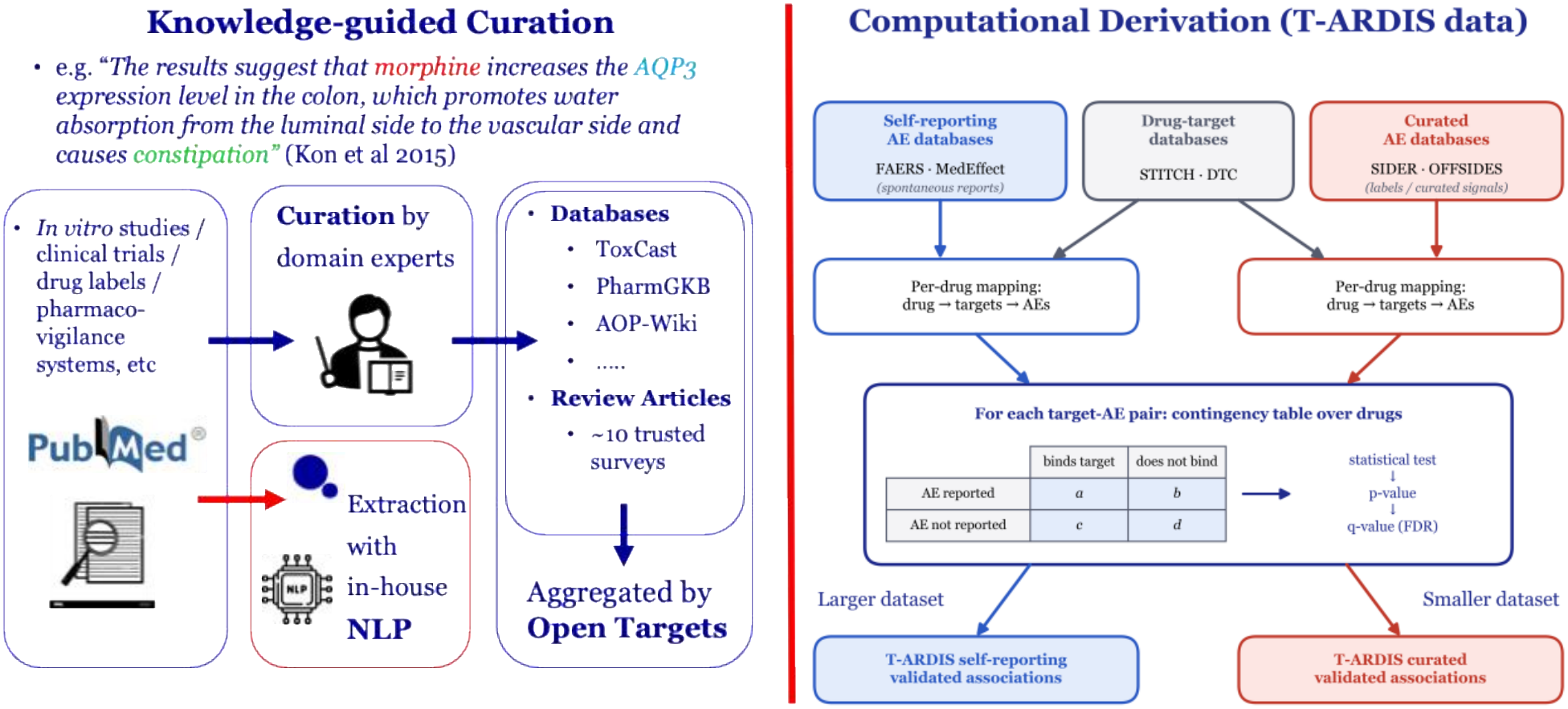
Pipeline used by Evotec to compile the ground truth target–AE associations in the TSA graph. Right panel redrawn after Galletti et al (2021).

### 2.3 Construction of the integrated TSA knowledge graph

The finalised TSA graph contains 11 node types and 20 edge types (Figures 2 and 3). Notably, of the 18,982 Gene nodes, only 473 (∼2.5%) appear in Gene_AE triples, reflecting the stringency of the false positive filtering policy and the limits of current target-AE knowledge.

**Figure 2:**
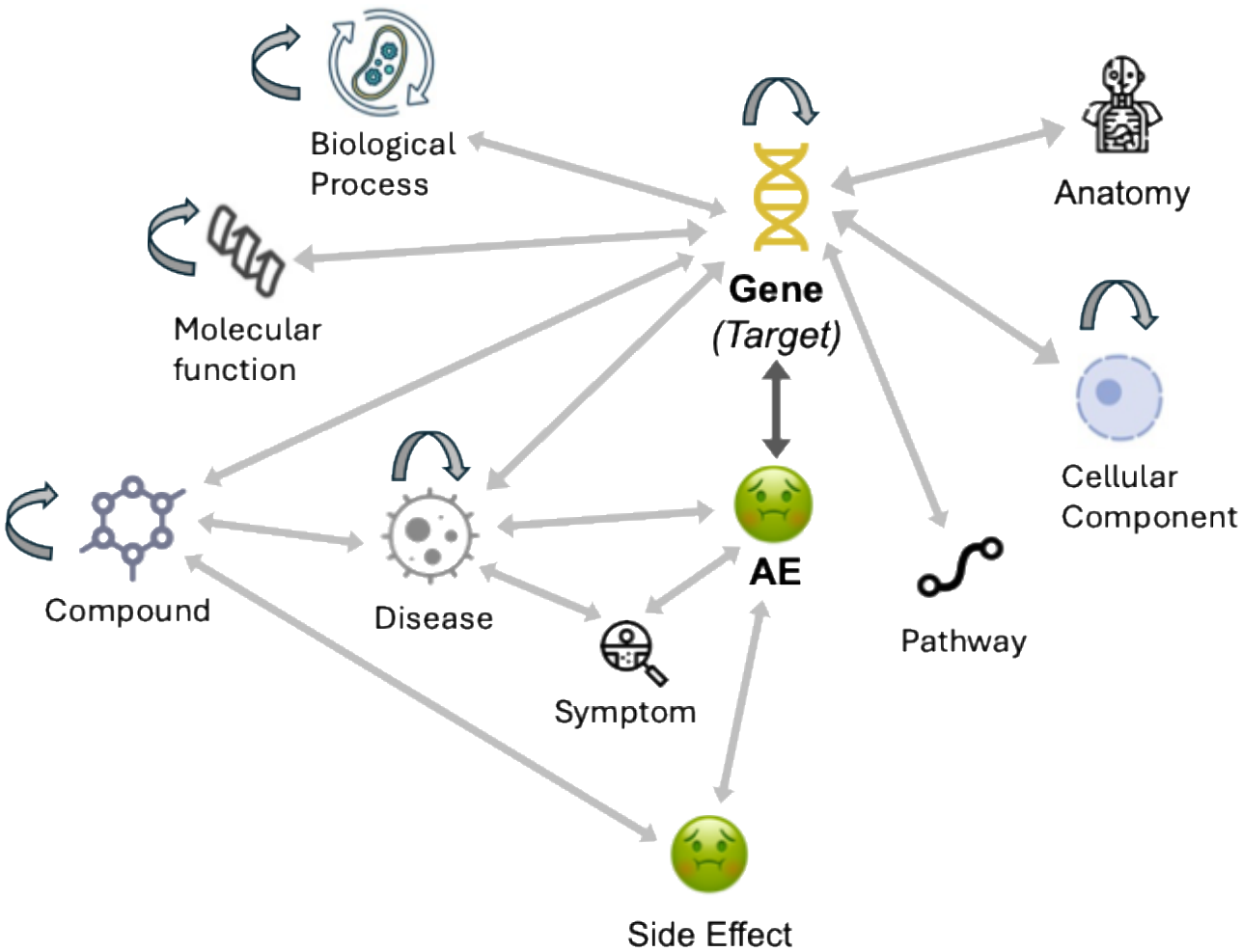
Schema of the TSA graph.

**Figure 3:**
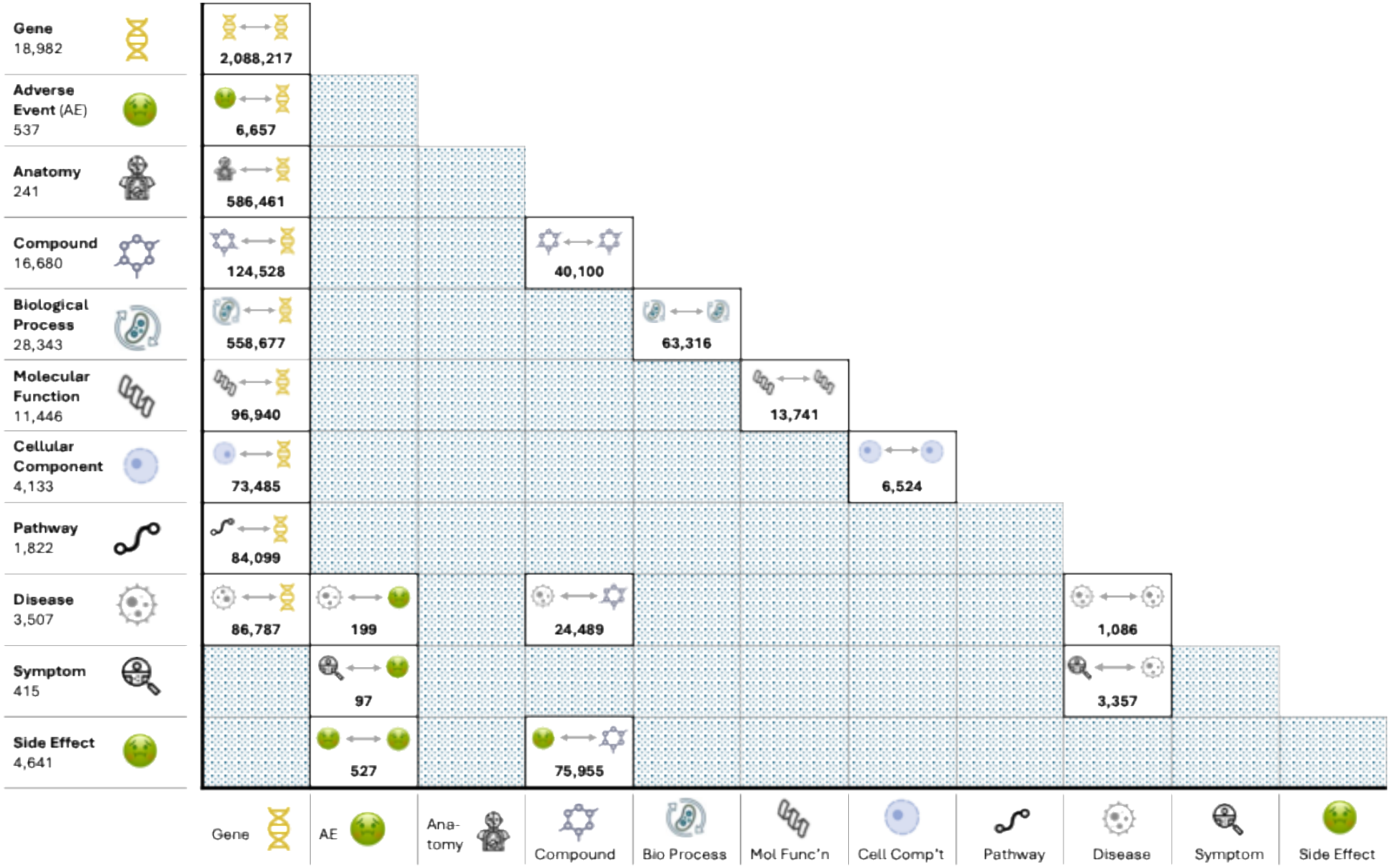
Node and edge type instances in the TSA graph. The vertical node index in the matrix shows the number of nodes of each type. The matrix’s cells show the number of instances of each edge type. Note that the TSA graph design enforces one edge type per node type pairing.

### 2.4 Topological bias in the TSA graph

An evaluation of the TSA graph revealed that topological imbalance strongly affects the distribution of target-AE associations. The top 20 targets by number of AE associations in the graph account for 2,997 target-AE associations, which is more than 45% of the total. Given this strong concentration of AEs among a small number of targets, target-AE association prediction on the TSA graph must integrate topological bias mitigation to be effective, thus motivating our main contribution. This is shown in Figure 4; for instance, drugs targeting CXCL8 can cause a wide variety of adverse events, including neutropenia and increased infection risk, because blocking this chemokine disrupts vital homeostatic neutrophil trafficking required to fight infectious diseases and manage systemic inflammation (Sitaru et al 2023, Russo et al 2014).

**Figure 4:**
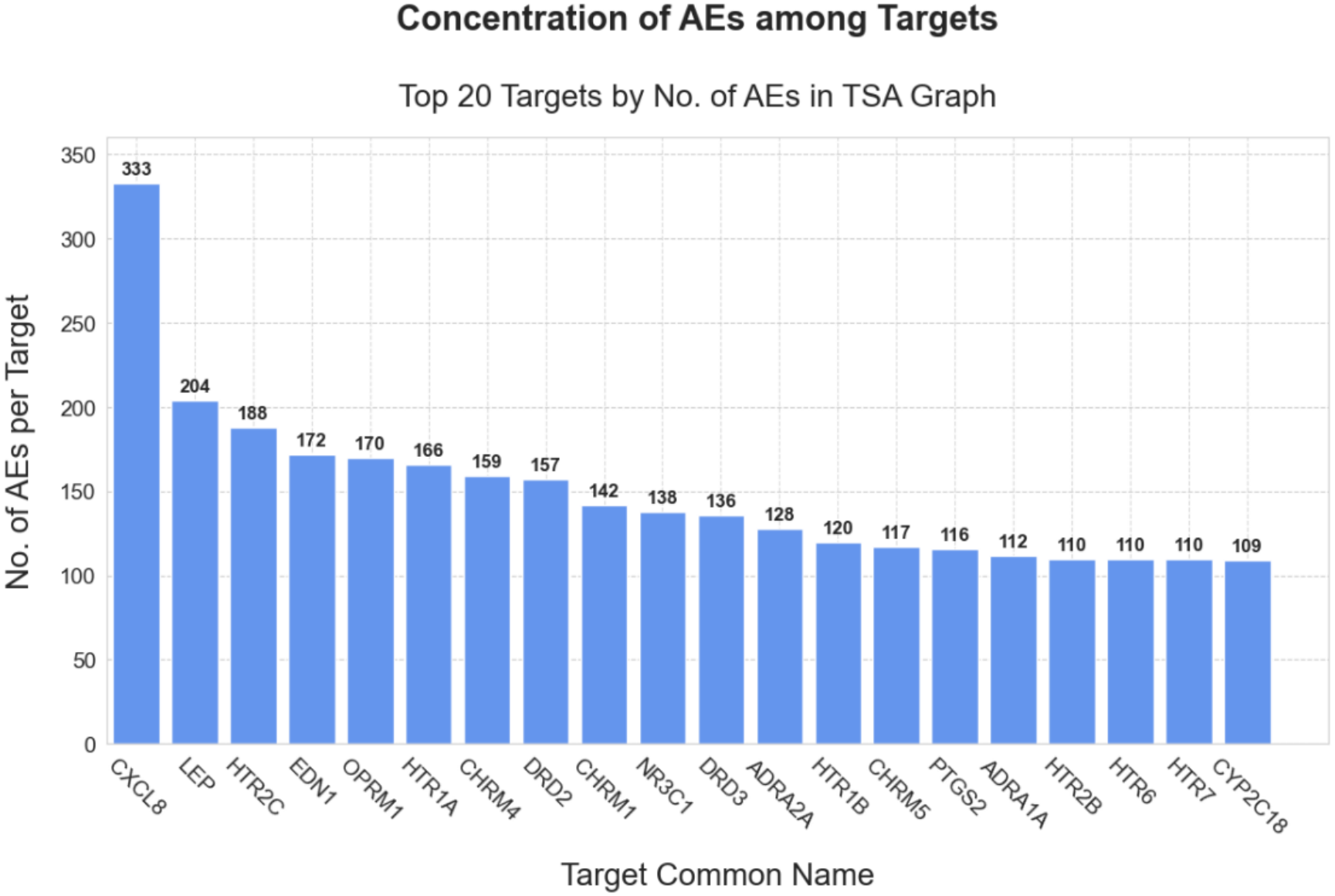
2,997 of the target-AE associations represent just 20 targets (denoted by their coding Genes in the TSA graph; common names taken from Entrez Gene). This amounts to ∼45% of the total number of associations. This strong level of concentration suggests a significant level of intrinsic “preferential attachment” – a natural tendency for a relatively small number of therapeutic targets to have a (putative) causal role in a large proportion of adverse events.

### 2.5 Target-AE association prediction as metapath-based KG completion

#### Defining the task

Our KG completion approach to target-AE association prediction represents a departure from prior efforts’ PPI-centric, single-AE models. The knowledge diversity achieved by integrating resources beyond PPIs allows the graph to encode potentially more effective predictive patterns. Furthermore, KG completion aims to train a model that can predict associated targets for all AEs in the graph, in contrast to the single-AE limitation of previous approaches.

The present work treats KG completion as a binary classification problem using KG derived predictive features. The 6,657 known target-AE associations whose curation is described in 2.2. form the positive class. A randomly selected subset of 15% (998) of these ground truth associations was set aside as a holdout test set. The remainder were used for 5-fold cross-validation and for training prior to testing,

To prevent data leakage during feature generation, any edges linking the entities of validation or test triples were permanently removed from the graph. This ensures that patterns containing the answers being predicted are not encoded in the features, preventing the model from learning on a graph that already includes the ground truth associations for which predictions are to be made.

Because non-existing target-AE associations are in practice not provable (and are indeed absent from reference databases), we treated the binary task as “positive against *assumed* negative”.

To facilitate model training under this condition, we employed a custom negative sampling policy. Examples were drawn from the pool of non-positive AE pairings of the 473 AE-associated targets (i.e. 473 ^*^ 537 - 6,657 = 254,001) to match the 5,659 non-holdout positives. We did not use targets with no known AE associations since we reasoned that these would tend to form trivially easy negatives for models to learn.

#### Evaluation approach

To better diagnose the unique performance challenges encountered when completing topologically biased KGs, we introduce a novel evaluation policy. Specifically, we evaluated models on validation data deciles representing different levels of existing target safety knowledge (e.g. deciles 1 and 2 represent targets with 0-5 AEs and 6-17 AEs, respectively). This allowed us to trace the impact of topological bias across the degree distribution of targets in the KG.

We used the true positive rate (TPR) in cross-validation for selecting model configurations to evaluate on test data. At the test stage, we supplemented TPR with the *hits@k* measure which provides insight into a KG completion model’s ability to rank known positive examples highly from among the remaining non-training pairings (henceforth “assumed negatives”).

In our case, ranking relies on the Random Forest’s positive class voting share as a continuous score by which to order positive and assumed negative entity pairings for the KG relationship in question (here target-AE associations). We accept the formulation of hits@*k* by Fey & Lenssen (2019), which computes it as the proportion of head entity + relation queries (i.e. targets with missing AEs) for which the correct tail entity (AE) ranks within the top *k* predictions from among all possible tail entities (all AEs in the graph). As the nature of target-AE associations permits multiple correct AEs per target, we employ *filtered hits@k* (Shi and Weninger 2017). This modification filters out higher-ranked correct AEs to ensure multiple true predictions do not demote each other. Throughout this work, we employ *k*=10.

#### Holdout Testing

Predictions on the holdout set were evaluated using (filtered) hits@10. We additionally report TPR. Confidence intervals for hits@10 and TPR were computed using bootstrap resampling.

#### External Testing

We utilised a set of target-AE associations compiled by Soldatos et al (2018) via an alternative methodology shown to be competitive with that by Kuhn et al (2013); to the best of our knowledge, our work presents the first use of this resource on the present task. We identified 7,804 associations from Soldatos et al’s collection that were unused assumed negative pairings in the TSA graph. We re-labelled these as positives for this testing scenario.

#### KG completion via metapath features

The goal of model interpretability, which is critical in the biomedical domain, steered our choice of KG completion classifier and features. All learned models are Random Forests (Breiman 2001), leveraging the clear link between feature values and prediction offered by the basic decision tree structure. We used KG-derived predictors known as “metapaths” as the model features by which to describe the two classes.

A metapath may be regarded as a “blueprint” for a path in the KG: it is a sequence of entity types linked by applicable relations. Thus, a given metapath may have multiple instances in a KG. In metapath-based KG completion approaches, We explored two distinct metapath feature sets: “L3”, which comprises 19 length-3 metapaths, and “L4”, an expanded set comprising 51 metapaths of lengths 3 and 4. The lengths offer a combination of path interpretability and predictive effectiveness (demonstrated in the domain beginning with Fu et al 2016) while maintaining comparatively low computational complexity. the frequencies of defined collections of metapaths are used as feature vectors to describe the KG triples formed by the entities at each end of the metapaths (Sun et al 2011).

### 2.6 Inverse Node Frequency (INF) feature transformation

INF translates the Inverse Document Frequency (IDF) metric (Spärck Jones 1972) to the knowledge graph context. INF replaces the raw frequency count for a metapath and is calculated as follows:

**Step 1:** Original or “raw” metapath feature computation

In the example in Figure 5, the original or raw frequency for the metapath (gene-compound-gene-AE) is 2 as there are two unique instances of this metapath for DCK–Rhinorrhoea.

**Figure 5:**
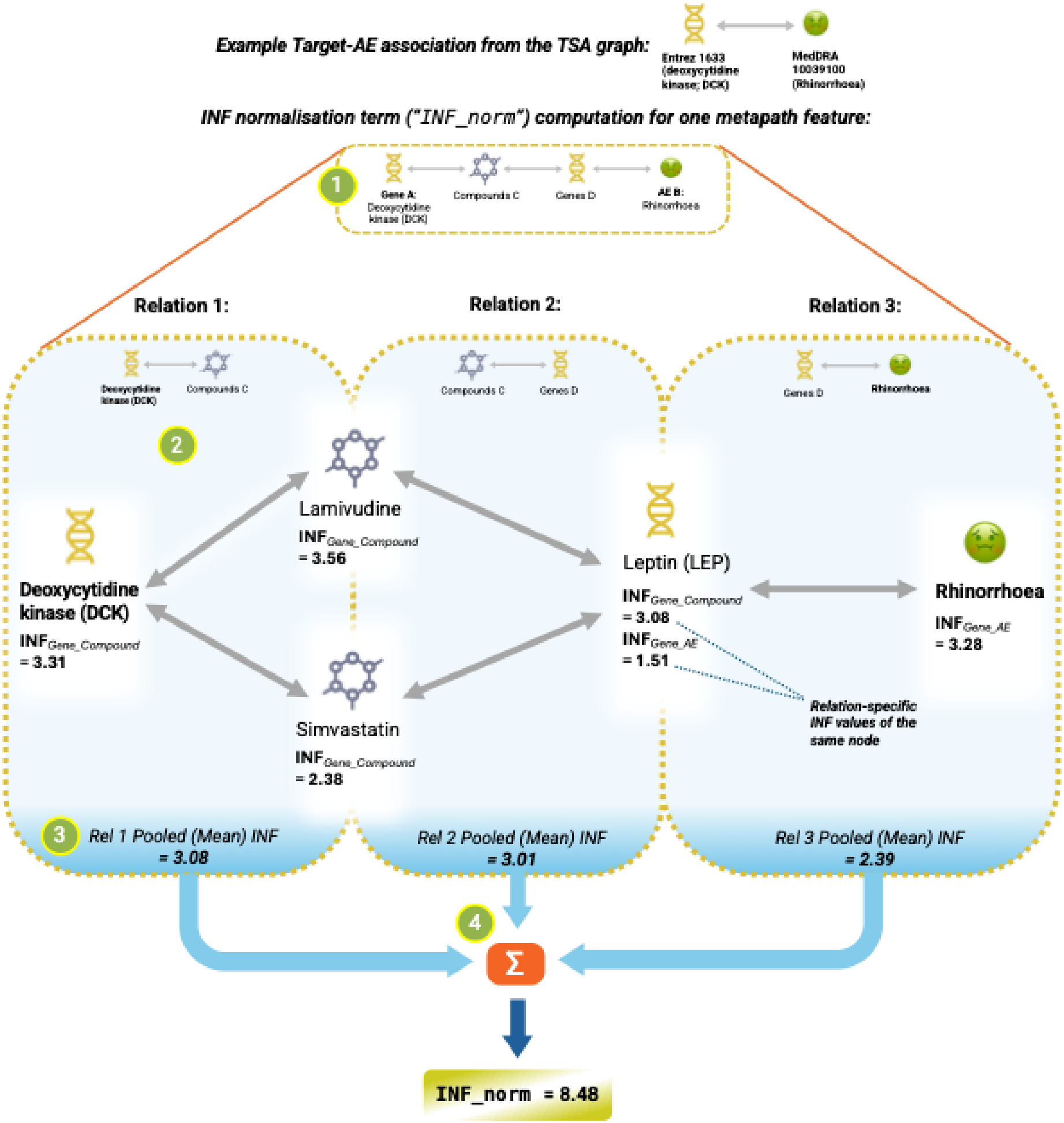
Computation of the INF normalisation term for a metapath feature of the association between the protein DCK and the AE Rhinorrhoea. The INF term is multiplied with the raw metapath feature value to complete the feature transformation.

**Step 2:** Individual-node INF computation

For a given node appearing in a given KG relation type, for example the DCK protein within the *Gene_Compound* relation in the TSA graph, as shown in Fig 5, the node’s INF value is calculated as:

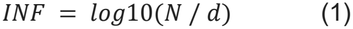

where *N* is the total number of triples with relation type *R* (i.e. *Gene_Compound*) in the graph, and *d* is the number of such triples that feature the node in question (i.e. where DCK is the *Gene* entity). For DCK in this context, its INF evaluates to 3.31 (to two decimal places).

**Step 3:** Path-Instance Weighting and Pooling

The INF value at each relation (or “hop”) of a metapath is determined by pooling the INF values of the nodes comprising that hop. In the example in Fig 5 these are the target DCK and both compounds linking to it. We evaluated three pooling operations: minimum, maximum, and mean. In the example, the mean-pooled INF at the first hop is (3.31+3.56+2.38)/3 = 3.08.

**Step 4:** Feature Aggregation and Deflation The relation-specific pooled INFs are then combined into a final INF normalisation term. We evaluated “sum”, “product”, and “None” as aggregation functions. Fig 5’s example illustrates the “sum” aggregation to arrive at 3.08+3.01+2.39 = 8.48 as the aggregated INF term. The original metapath count is multiplied by this INF term to arrive at the INF-transformed metapath feature value.

To further control the influence of extreme raw counts caused by high-degree nodes, a deflator exponent was introduced as an additional parameter (values tested: 0, 0.3, 0.5, 0.75, 1). This is applied to the metapath count prior to applying the INF term.

The set of INF transformation parameters, their functions and the values that were explored in cross-validation are presented in Table 1. The resulting set of 35 unique parameter configurations is included in the Supporting Information.

**Table 1.**
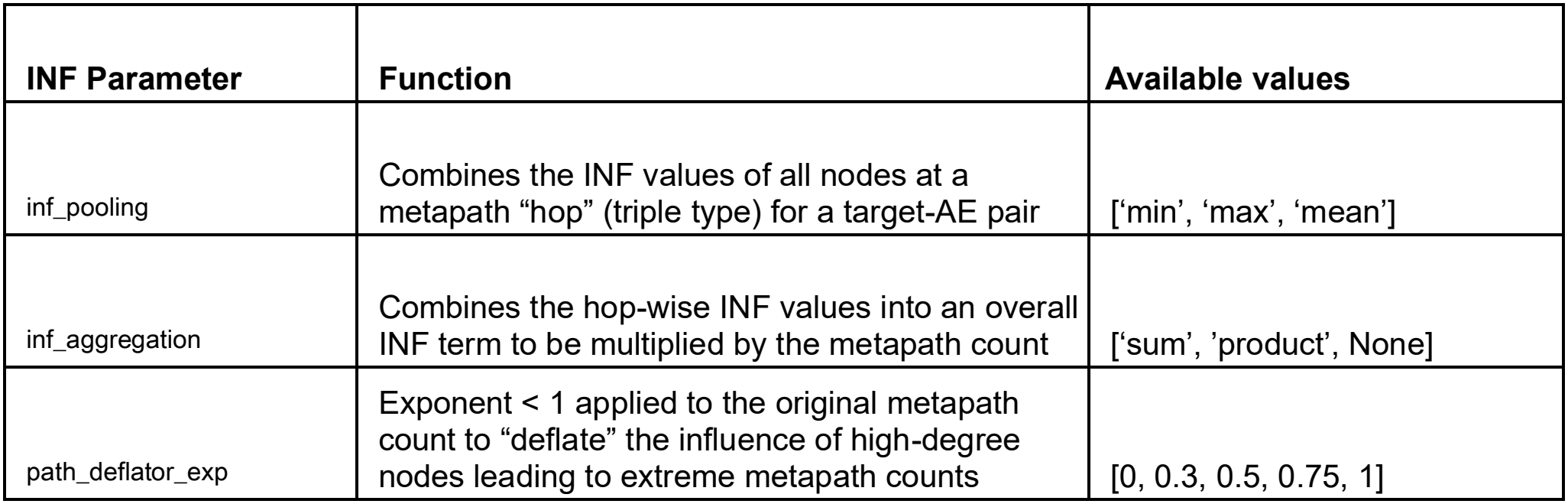
INF transformation parameters and their values explored in the present study.

### 2.7 Benchmarking

To establish a baseline and assess the impact of topological imbalance, models were initially trained using the original, untransformed metapath count features.

As an alternative transformation for tackling topological bias, we selected the Degree-Weighted Path Count (DWPC) introduced by Himmelstein et al (2015, 2017). DWPC was designed to reduce the influence of high-degree hub nodes on metapath feature values.

In DWPC each metapath instance contributes the product of its constituent nodes’ down-weighted degrees (using a negative exponent; -0.4 per Himmelstein et al 2017); the transformed metapath feature value is the sum of these contributions. DWPC has been utilised in recent drug discovery applications (Binder et al 2022; Ranjbar et al 2023).

#### Hyperparameter Optimisation

All Random Forest classifiers were trained with 200 trees using scikit-learn (Pedregosa et al 2011); hyperparameters were explored via grid search during cross-validation. See the Supplementary Information for further model implementation details.

## 3 Results

### 3.1 Cross-validation confirms topological imbalance degrades prediction for sparsely annotated targets

Figure 6 visualises TPR on three selected validation deciles by number of known AE associations per target (as defined in 2.5). These comprise the 1^st^ decile (associations where targets have 0-5 known AEs), the 2^nd^ decile (targets with 6-17 AEs) and the 10th decile (targets with 163-256 AEs). The untransformed metapath features achieved near-perfect TPR on the 10^th^ decile: 0.96 for L3 features and 0.97 for L4 features). Conversely, performance collapsed on the 1^st^ decile, with TPR at only 0.20 for L3 features and 0.22 for L4 features. These results confirmed that topological imbalance severely degrades performance on targets with sparse safety profiles and motivated the exploration of the proposed INF transformation on this task.

**Figure 6:**
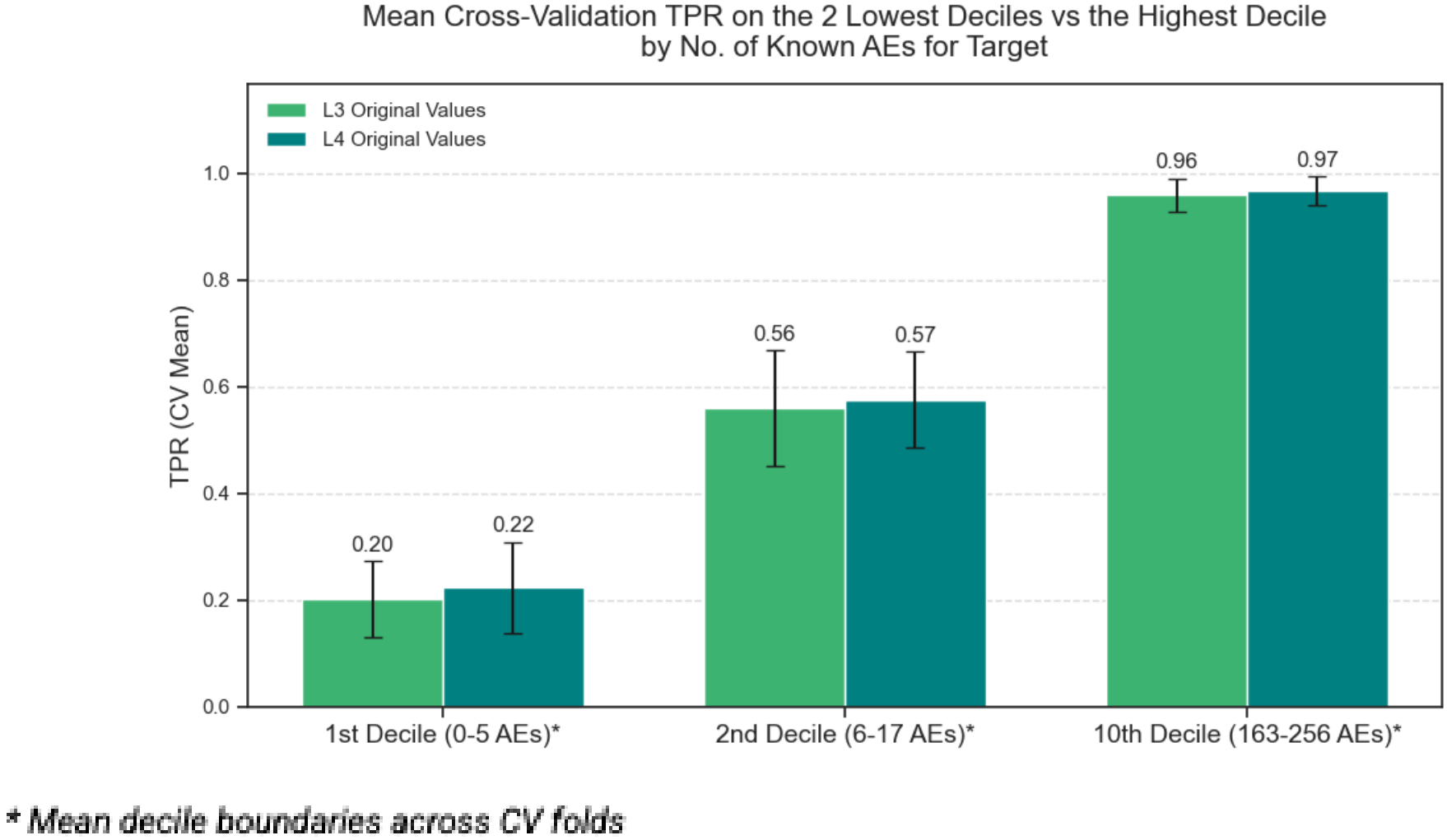
Bar chart illustrating the contrast in CV TPR between the two lowest deciles and the single highest decile of validation data by no. of known AEs for the target. Error bars are t-based 95% confidence intervals on the CV TPR mean across the 5 cross-validation folds.

Furthermore, the additional 32 metapath features in the L4 feature set compared to L3 clearly offered no meaningful performance gains, despite requiring substantially more computational time for generation (∼10 hours for L4 versus ∼3 hours for L3). Consequently, evaluations on test data proceeded utilising only the L3 feature set.

### 3.2 INF transformation mitigates degree bias and improves predictions for low-degree targets

To determine if the proposed Inverse Node Frequency (INF) transformation successfully mitigates degree bias, we compared the CV TPR of models utilising INF-transformed features against models using the original features and the Degree-Weighted Path Count (DWPC) benchmark. Across all validation data, the top three INF configurations achieved higher mean CV TPR (up to 0.87) than both the DWPC (0.85) and original (0.84) benchmarks. The best-performing general INF setting utilised ‘mean’ pooling, ‘product’ aggregation, and a 0.3 deflator exponent.

The superiority of the INF transformation was most pronounced in the challenging lowest-data decile (Figure 7). On this data subset (target-AE associations where targets have 0-5 known AEs), the best INF configuration achieved a TPR of 0.42, more than doubling the performance of the original features (0.20) and greatly outperforming DWPC (0.26).

**Figure 7:**
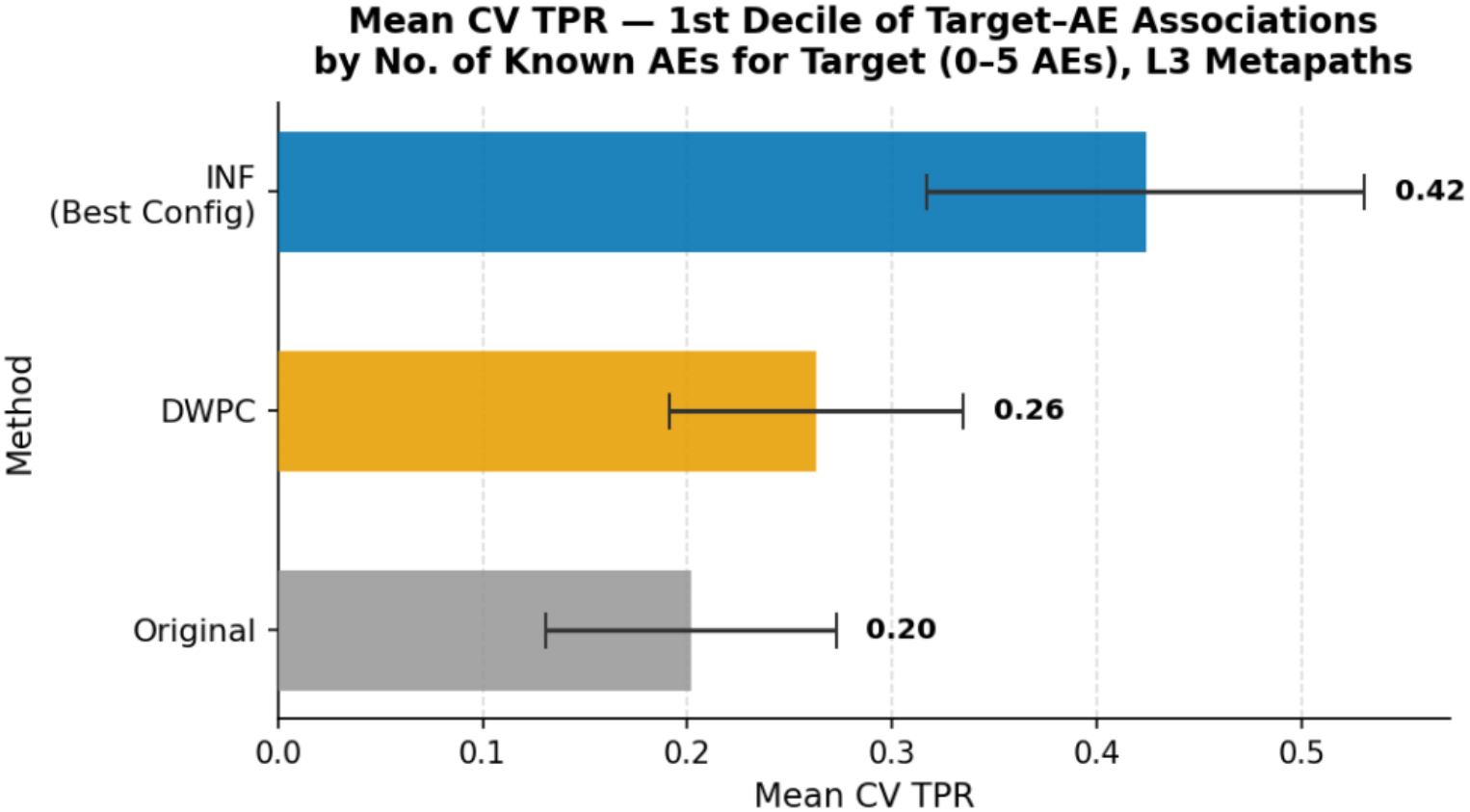
The INF transformation shows superiority on the lowest decile of validation data when using L3 features. Error bars are t-based 95% confidence intervals on the CV TPR mean across the 5 cross-validation folds.

Notably, the top three INF configurations for this specific decile (parameters shown in Table 2) all utilised a deflator exponent of 0: the original metapath value (i.e. the count of instances of that metapath) is completely deflated away by applying an exponent of 0 (INF). This indicates that the original metapath values are unimportant for TPR on this distinct, challenging subset of the validation data. As all three configurations do include an INF term, the result suggests that the INF transformation specifically benefits performance on this data decile.

**Table 2.** Parameter values of the three best INF configurations by mean CV TPR on the 1^st^ decile.

| INF configuration (rank) | INF parameter values |  |  |
|---|---|---|---|
|  | inf_pooling | inf_aggregation | path_deflator_exp |
| 1 | ‘max’ | ‘product’ | 0 |
| 2 | ‘mean’ | ‘product’ | 0 |
| 3 | ‘max’ | ‘sum’ | 0 |

Motivated by the improvement on the 1^st^ decile, we constructed a decile-best INF ensemble: for each target-AE candidate pair the ensemble returns the prediction from the single INF configuration that had performed best on that target’s CV decile. The ensemble achieved an overall mean CV TPR of 0.89, whereas the single best overall configuration achieved 0.87.

Results across all 35 configurations showed that the best settings on the two lowest deciles all suppress the raw count entirely (deflator exponent = 0, i.e. prediction by the INF term alone), while at higher deciles no single deflator exponent dominates.

Overall and per-decile CV results across all 35 configurations are reported in Table S1. Table S2 lists the top 3 INF configurations for each decile. Figure S3 is an additional visual aid for comparing the CV TPR of the single best overall configuration and the decile-best configuration for each decile, with raw-count results also depicted.

### 3.3 Holdout test set evaluation reveals divergence between ranking and classification metrics

When evaluating the entire holdout set via the ranking-based hits@10 metric, all models proved highly competitive and performed similarly, with DWPC and the original features at 0.36, closely followed by the INF ensemble (0.35). However, the classification-based TPR metric revealed a different dynamic (Supplementary Figure S4): the INF ensemble dominated with a TPR of 0.90, outperforming DWPC (0.87) and the original features (0.85).

This divergence between hits@10 and TPR was especially clear when isolating the holdout test data involving the lowest-degree targets (0-5 known AEs; Figure 8). For hits@10 on this challenging subset, the INF ensemble closely matched DWPC and outperformed the original features. However, when evaluating TPR on this same subset, the INF ensemble correctly identified nearly twice as many true positives (0.48 TPR) as the DWPC method (0.26) and the original features (0.20). This suggests that while ranking true positives tightly into the top 10 is difficult across all methods for low-degree targets, the INF transformation significantly improves the broader recovery of these challenging biological associations. In practice, modellers would thus be well advised to take into consideration all target-AE associations predicted by INF-transformed features rather than focusing on a narrow top *k* ranking.

**Figure 8:**
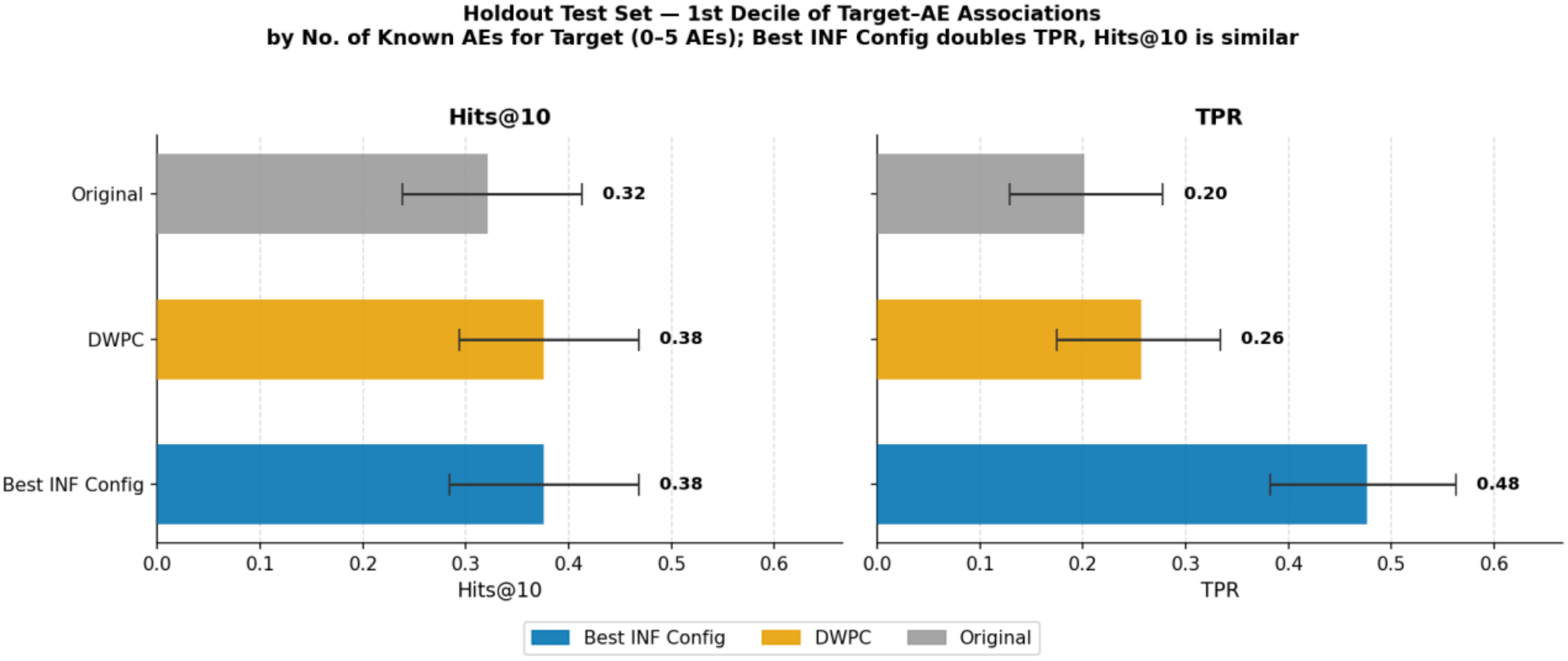
Holdout test set performance on the 1st decile of target-AE associations (targets with 0-5 known AEs, n = 109 associations). Left panel: (filtered) Hits@10; right panel: TPR (threshold 0.5). Both panels share the same y-axis scale. The best INF configuration (selected by 1st-decile CV TPR) doubles TPR relative to DWPC and the raw-count baseline, while Hits@10 is broadly comparable across methods. Error bars are 95% bootstrap confidence intervals (1,000 resamples).

### 3.4 External validation demonstrates the generalisability of INF-transformed features

To assess the generalisability of the learned patterns, the models were challenged with an external test set of 7,804 target-AE associations matched from the collection published by Soldatos et al (2018). Figure 9 contrasts model performance in terms of hits@10 and TPR on this external resource. Hits@10 scores dropped sharply for all models (falling below 0.07), strongly indicating that the hits@10-relevant predictive patterns learned from the graph were specific to the original ground truth data distribution.

**Figure 9:**
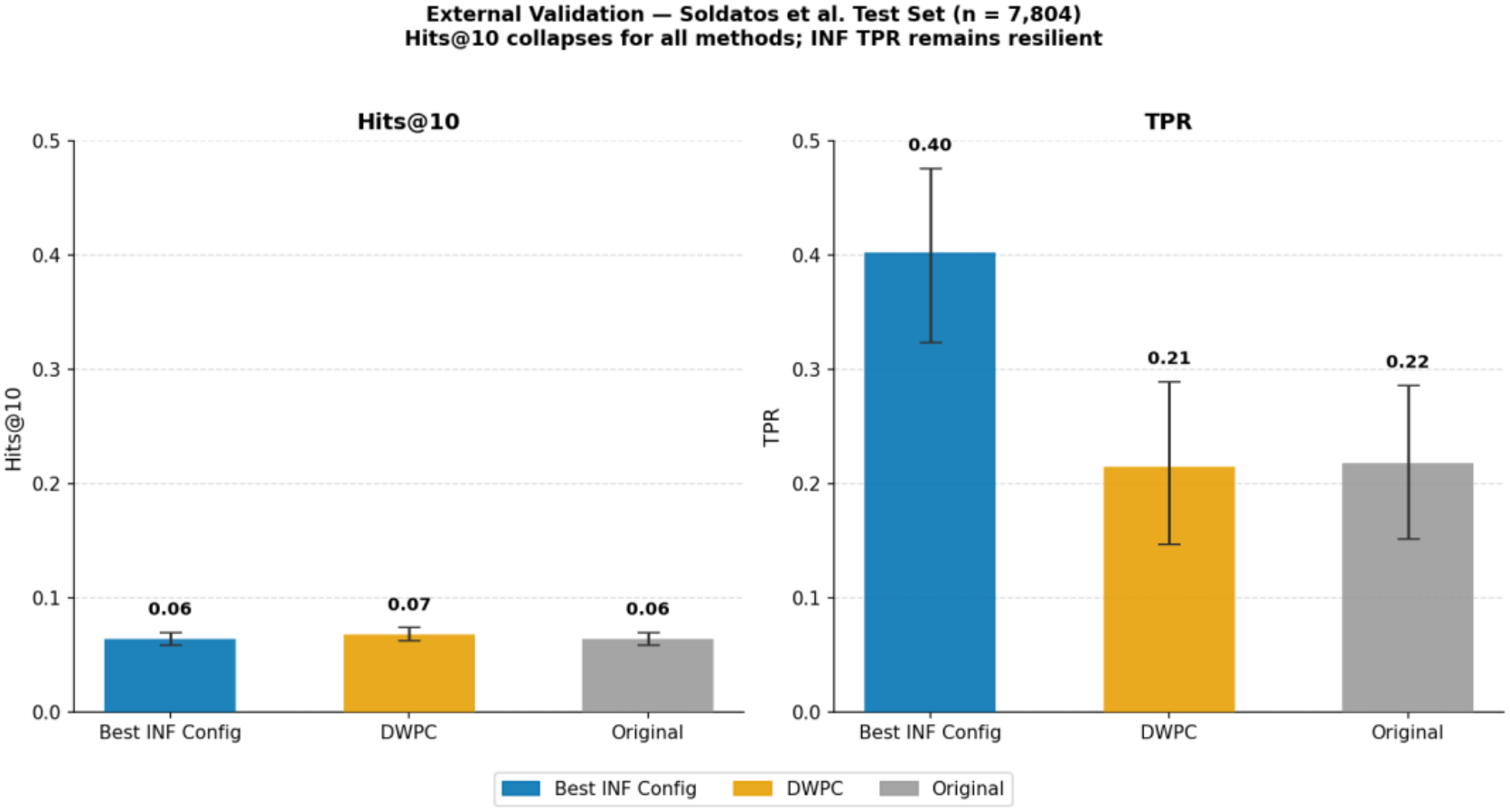
External validation on the Soldatos et al (2018) dataset (n = 7,804 target–AE pairs). Left panel: filtered Hits@10; right panel: TPR. Both panels share the same 0-0.5 y-axis scale. Error bars are 95% bootstrap confidence intervals (1,000 resamples).

Despite the collapse in ranking performance, the INF ensemble demonstrated visible resilience in classification performance compared to the alternatives. The INF ensemble maintained an overall TPR of 0.40 on the external dataset, nearly double the TPR achieved by the original features and DWPC (both at 0.22). This resilience was driven mostly by performance on the low-data associations, which made up the majority (5,183 out of 7,804) of the external test set. In the 1st decile subset (0-5 AEs) of the external data, the benchmarks achieved negligible TPRs (< 0.05), whereas the INF ensemble maintained a comparatively robust TPR of 0.25. This result is directly relevant to target safety assessment in drug discovery: retaining recall on external associations means the INF-based model can support safety profiling for targets beyond the original ground truth, including for understudied targets that motivate computational TSA in the first place.

### 3.5 Feature importance analysis highlights the mechanistic value of drug-target co-occurrence

To determine which portions of the knowledge graph were driving predictions, we utilised permutation importance analysis on the subset of holdout and external test data comprising targets with 0-5 AEs (Figures 10 and 11). There was notable overlap among the top five metapath features driving performance for both the INF model and the benchmarks.

**Figure 10:**
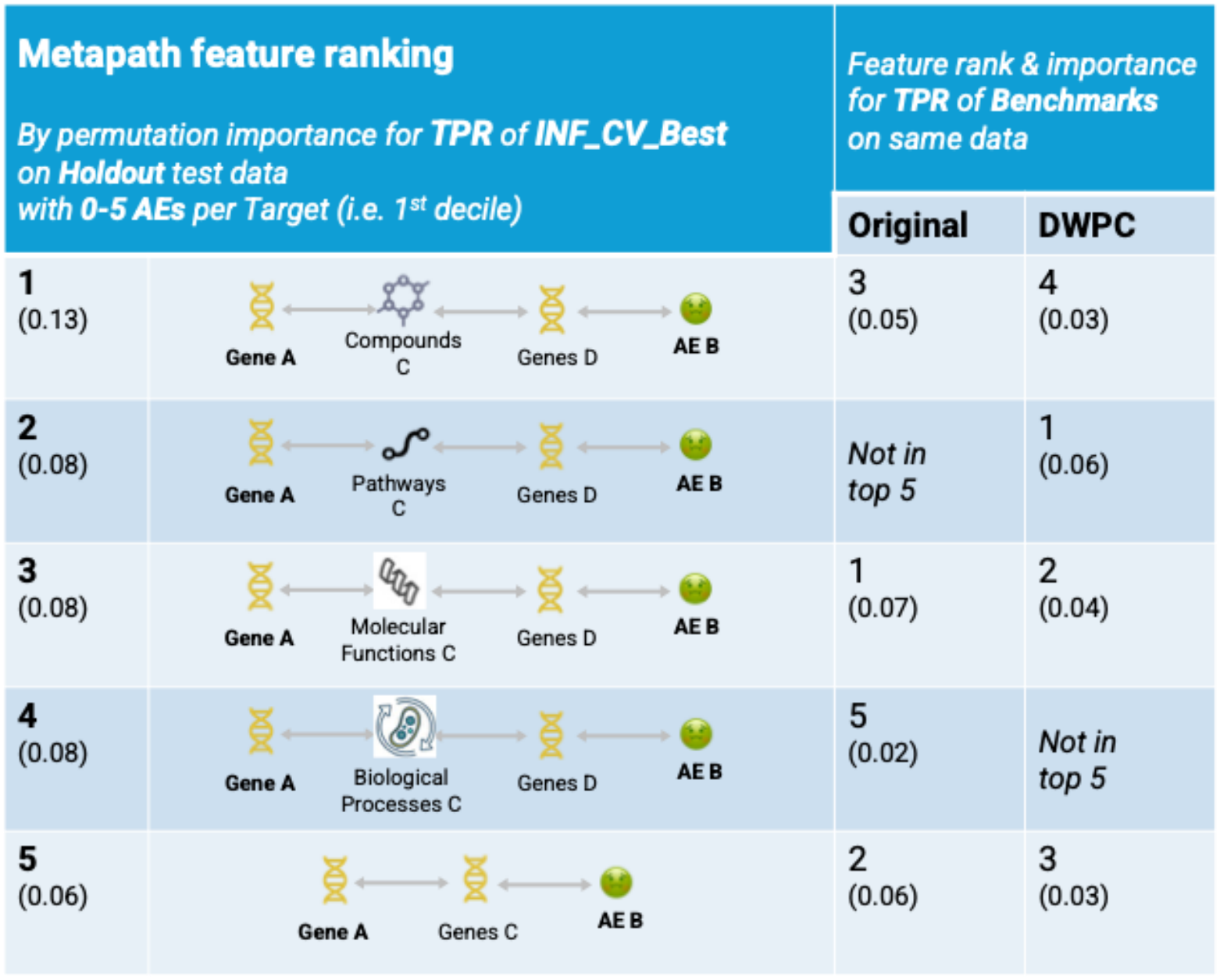
Top five features for TPR of the INF model on holdout test data with 0-5 AEs per target (named “INF_CV_Best” as it produced the highest TPR for such data during cross-validation).

**Figure 11:**
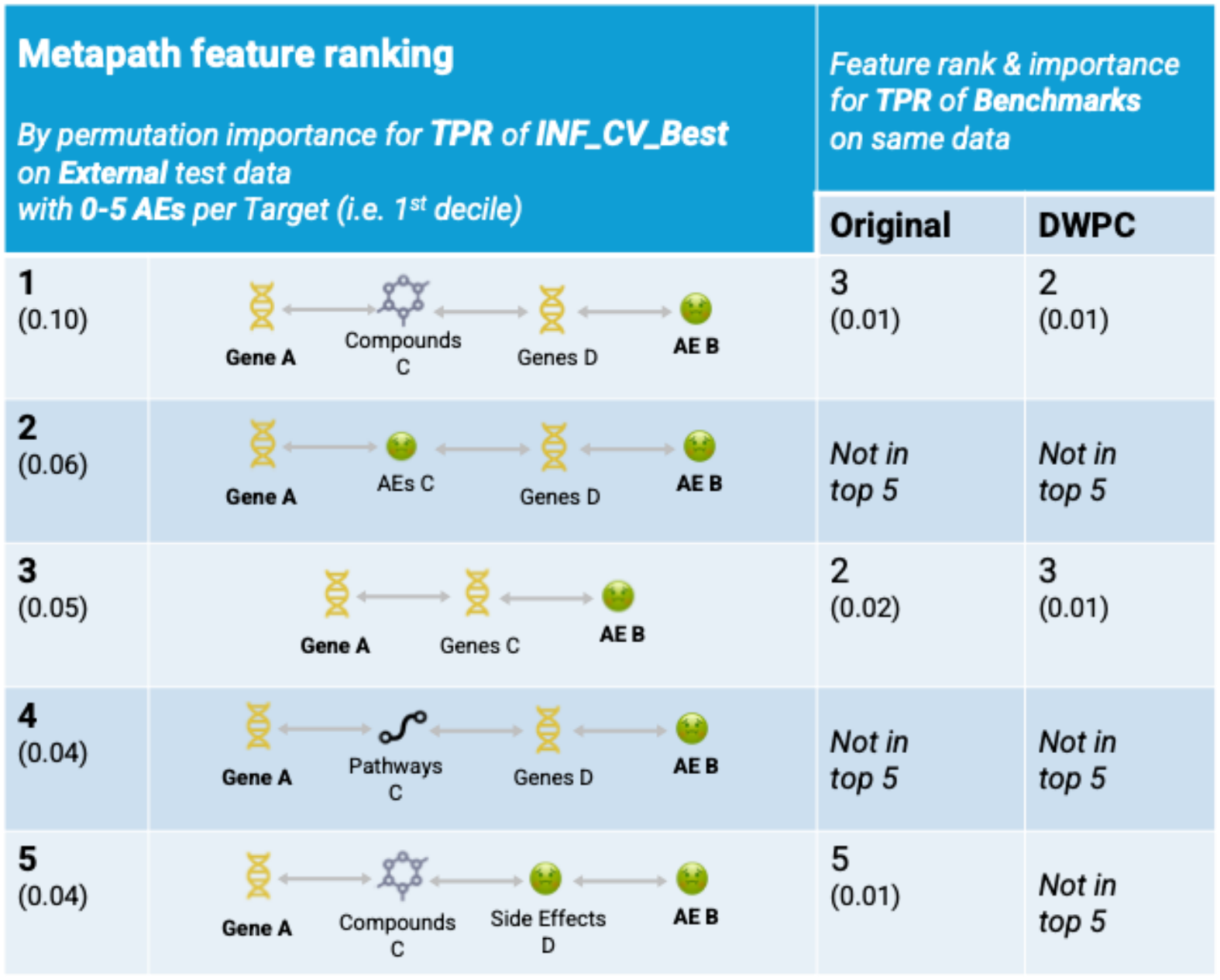
Top five features for TPR of the INF model on external test data (Soldatos et al 2018) with 0-5 AEs per target (named “INF_CV_Best” as it produced the highest TPR for such data during cross-validation).

Notably, the single most important feature for the INF model across both hits@10 and TPR was the metapath representing Gene_Compound_Gene_AE (signalling that the target and AE are linked via shared compounds and other targets). This relies on the co-occurrence of drug-target and AE combinations, aligning with the premise of statistical pharmacovigilance approaches (such as those by Kuhn et al and Soldatos et al) and the biological premise that proteins that share ligands often share biological properties (Keiser et al 2007).

Furthermore, on the external test set, the INF model continued to rank this mechanistic metapath as its most important feature, whereas the benchmarks struggled to assign significant importance to any individual metapaths, further supporting INF’s ability to prioritise relevant domain knowledge.

## 4 Discussion

The primary reason why target-AE association prediction via KG completion is challenging is that low-data entities have sparse topological profiles in biomedical KGs. Their AE associations tend to lie outside the training distribution of any model that learns to associate target connectivity with AE association likelihood. We have shown that KG completion with untransformed metapath features demonstrates this weakness; prior work by Bonner et al (2022a) indicates that popular KG embedding methods are equally vulnerable.

The reference correction for this bias in the metapath literature, the Degree-Weighted Path Count (DWPC), is only a partial fix. DWPC normalises a metapath’s count using the absolute degrees of its constituent nodes. It therefore corrects for “bulk” connectivity, without distinguishing relation-specific topological profile, placing an inherent limit on the expressivity of the normalisation.

INF is explicitly relation-specific by design, with each KG relation type contributing equitably to the transformation via the pooling and aggregation steps described in 2.6. The results in 3.2 show that the best-performing INF configurations on the two deciles with the fewest known AEs per target are precisely those that suppress the raw metapath count entirely, with predictions driven by the INF term alone. Under these configurations, metapath feature value is greatest when all entities in the traversed edges are sparsely connected, potentially suggesting biologically specific association, in contrast to the noisy connectivity of hub entities. The first recommendation flowing from these findings is direct: for target safety assessment applications, the INF transformation should be preferred over both the untransformed metapath count baseline and the established DWPC correction. The recall gains observed at the lowest deciles translate directly into improved performance on the understudied targets that constitute the primary use case for computational TSA, and the advantage holds under the external distribution shift represented by the Soldatos et al (2018) test set.

A pattern was observed in our results wherein INF was competitive against both the raw metapath counts and the DWPC normalisation on holdout and external data Hits@10, but substantially stronger on TPR, particularly at the lowest decile (associations with 0-5 AEs per target). This suggests that the INF transformation in its current form raises model scores close to the class threshold (thus raising TPR) but may also depress scores for some high-confidence cases, occasionally pushing them out of the top k. The implication is that, for INF and methods like it, CV TPR is a reliable predictor of test data TPR but not of test data Hits@k. Beyond the practical recommendation above, the divergence between the two metrics motivates innovation in ranking-based evaluation. As noted by Bonner et al (2022b), a strict focus on the top k may obscure valuable lower-ranked predictions. Procedures that characterise the distribution of true positives across the full rank order could more faithfully capture model behaviour. Such innovations would also transfer to other prioritisation problems on biomedical KGs, such as disease-associated gene prediction.

One limitation of our work is that training and holdout evaluation rely on the single curated ground truth set compiled at Evotec. Although the external validation on a subset of the collection by Soldatos et al (2018) provides an initial test of generalisation, each new practical applications is likely to reveal its own unique distributional shift. It’s clear from the results of the external validation set that training data in target safety evaluation is small and models are prone to overfitting probably because of the limited examples the models can learn from. Furthermore, the inherent uncertainty around assumed negatives is not addressed by INF.

## 5 Conclusion and follow-on work

This study has explored target-AE association prediction as a KG completion problem. Topological imbalance in the underlying KG was found to severely skew performance by raw metapath count features, favouring associations involving targets with substantial existing safety knowledge and producing a high false negative rate on targets with sparser safety profiles. We introduced Inverse Node Frequency, a metapath feature transformation that recovers signal from these low-data associations by weighting each path instance by the inverse frequency of its intermediate nodes within their distinct relation types. INF displayed robust competitiveness throughout cross-validation, and the configurations recommended by CV exhibited clear TPR resilience on external test data.

INF can be adapted and generalised beyond metapath-based target-AE associations prediction. INF is a flexible graph topology transformation rather than a prescribed model architecture; any KG completion approach which depends on entity connectivity can be modified with a relation-specific rarity term. We see at least two concrete insertion points for INF in other popular KG completion methods.

In KG embedding-based methods, a node’s INF values across all relation types in the given KG can be arranged into a vector and supplied alongside the learned embedding as an additional hand-crafted feature or used as a prior upon embedding initialisation.

In KG-adapted graph neural network (GNN) methods like R-GCN (Schlichtkrull et al 2018), each element in a KG relation’s adjacency matrix can be weighted by the INF of one or both entities of the corresponding connected pair. In this manner, the characteristic GNN aggregation over the node neighbourhood will be automatically biased away from high-degree neighbours and towards sparsely connected ones likely to encode specific roles in the network.

INF’s apparent ability to surface paths whose rarity may carry domain-grounded meaning makes it a natural candidate for interdisciplinary follow-up: validating, with input from subject matter experts, whether the patterns INF prioritises encode genuine biological specificity across diverse biomedical KG completion problems. Refining these practices can clear a new path to trustworthy safety assessment for the targets where it is most needed.

## Supporting information

Supplementary Information Pack

## Acknowledgements

The authors thank the *In Silico R&D* team at Evotec for their contribution to the curation of the target-adverse event associations.

The AI assistant Claude (Anthropic), used via Claude Code, assisted with: (i) cleaning, documenting and testing the analysis code, and writing the figure-generation scripts and automated reproduction tests in the accompanying repository; (ii) checking figure and section cross-references. The AI tool did not design or draft the study, the INF method or its code implementation, run the analyses or interpret the results. The authors reviewed, edited and verified all AI-assisted code and text, and take full responsibility for the content of this article.

## Data Availability

The code implementing INF, the reproduction notebook, tests and the derived result files needed to reproduce the figures are freely available under the MIT License at https://github.com/TERENTIVS/INF_KGC; the submitted version is archived at Software Heritage (swh:1:rev:09ff3525acb878c37a58d744b32ff11e45b14fad). The TSA knowledge graph was derived from the publicly available Drug Repurposing Knowledge Graph (Ioannidis et al 2020). The external validation data are from Soldatos et al (2018) and are redistributed under their original CC BY 4.0 licence. The curated target-adverse event associations used as ground truth are proprietary to Evotec and cannot be shared publicly. Pre-computed features and predictions that contain these associations, and a sample documenting the data schema, are available on reasonable request from the first author (T.M. Egbelo). All values reported in the figures are provided in the repository and can be verified without the restricted data.

