## Supplementary Information Pack for "Improving Target–Adverse Event Association Prediction via Inverse Node Frequency Transformation to Mitigate Knowledge Graph Topological Bias"

**Table S1:** Cross-validation TPR (recall) for all 35 INF transformation configurations (rows), at each decile of target safety knowledge (columns); * marks each decile-best INF configuration.

| Configuration | D1 | D2 | D3 | D4 | D5 | D6 | D7 | D8 | D9 | D10 | Overall |
|---|---|---|---|---|---|---|---|---|---|---|---|
| 0 \| raw counts (baseline) | 0.20 | 0.56 | 0.87 | 0.97 | 0.98 | 0.99 | 0.97 | 0.98 | 0.99 | 0.96 | 0.85 |
| 1 \| defl=–, agg=sum, pool=min | 0.23 | 0.61 | 0.89 | 0.97 | 0.98 | 0.99 | 0.97 | 0.98 * | 0.98 | 0.96 | 0.86 |
| 2 \| defl=–, agg=sum, pool=max | 0.23 | 0.63 | 0.88 | 0.97 | 0.99 | 0.99 | 0.97 | 0.98 | 0.99 | 0.96 | 0.86 |
| 3 \| defl=–, agg=sum, pool=mean | 0.23 | 0.61 | 0.88 | 0.97 | 0.99 * | 0.99 | 0.97 | 0.98 | 0.98 | 0.96 | 0.86 |
| 4 \| defl=–, agg=product, pool=min | 0.25 | 0.62 | 0.89 | 0.98 * | 0.99 | 0.99 | 0.97 | 0.98 | 0.99 | 0.97 | 0.86 |
| 5 \| defl=–, agg=product, pool=max | 0.25 | 0.62 | 0.88 | 0.97 | 0.98 | 1.00 * | 0.97 | 0.98 | 0.99 * | 0.97 | 0.86 |
| 6 \| defl=–, agg=product, pool=mean | 0.26 | 0.62 | 0.88 | 0.98 | 0.98 | 0.99 | 0.97 | 0.97 | 0.99 | 0.96 | 0.86 |
| 7 \| defl=0, agg=–, pool=– (deflated count only) | 0.36 | 0.66 * | 0.80 | 0.85 | 0.77 | 0.91 | 0.91 | 0.94 | 0.86 | 0.84 | 0.79 |
| 8 \| defl=0, agg=sum, pool=min | 0.21 | 0.45 | 0.80 | 0.96 | 0.96 | 0.99 | 0.97 | 0.98 | 0.98 | 0.96 | 0.83 |
| 9 \| defl=0, agg=sum, pool=max | 0.40 | 0.58 | 0.79 | 0.87 | 0.87 | 0.94 | 0.95 | 0.96 | 0.94 | 0.89 | 0.82 |
| 10 \| defl=0, agg=sum, pool=mean | 0.40 | 0.57 | 0.76 | 0.90 | 0.88 | 0.96 | 0.92 | 0.96 | 0.90 | 0.87 | 0.81 |
| 11 \| defl=0, agg=product, pool=min | 0.22 | 0.47 | 0.80 | 0.93 | 0.95 | 0.98 | 0.97 | 0.98 | 0.98 | 0.96 | 0.83 |
| 12 \| defl=0, agg=product, pool=max | 0.42 * | 0.60 | 0.77 | 0.90 | 0.86 | 0.94 | 0.95 | 0.97 | 0.92 | 0.90 | 0.82 |
| 13 \| defl=0, agg=product, pool=mean | 0.42 | 0.61 | 0.77 | 0.90 | 0.87 | 0.94 | 0.94 | 0.96 | 0.88 | 0.88 | 0.82 |
| 14 \| defl=0.3, agg=–, pool=– (deflated count only) | 0.21 | 0.55 | 0.88 | 0.98 | 0.98 | 0.99 | 0.97 | 0.98 | 0.99 | 0.96 | 0.85 |
| 15 \| defl=0.3, agg=sum, pool=min | 0.25 | 0.64 | 0.91 * | 0.97 | 0.98 | 0.99 | 0.97 | 0.98 | 0.98 | 0.97 | 0.86 |
| 16 \| defl=0.3, agg=sum, pool=max | 0.25 | 0.59 | 0.88 | 0.98 | 0.98 | 1.00 | 0.97 | 0.98 | 0.99 | 0.97 | 0.86 |
| 17 \| defl=0.3, agg=sum, pool=mean | 0.25 | 0.61 | 0.88 | 0.98 | 0.99 | 0.99 | 0.98 * | 0.97 | 0.99 | 0.97 | 0.86 |
| 18 \| defl=0.3, agg=product, pool=min | 0.31 | 0.55 | 0.87 | 0.96 | 0.95 | 0.99 | 0.97 | 0.97 | 0.96 | 0.96 | 0.85 |
| 19 \| defl=0.3, agg=product, pool=max | 0.29 | 0.59 | 0.87 | 0.97 | 0.98 | 1.00 | 0.97 | 0.98 | 0.99 | 0.97 | 0.86 |
| 20 \| defl=0.3, agg=product, pool=mean | 0.33 | 0.63 | 0.89 | 0.98 | 0.97 | 1.00 | 0.98 | 0.98 | 0.99 | 0.97 * | 0.87 * |
| 21 \| defl=0.5, agg=–, pool=– (deflated count only) | 0.21 | 0.55 | 0.88 | 0.98 | 0.98 | 0.99 | 0.97 | 0.98 | 0.99 | 0.96 | 0.85 |
| 22 \| defl=0.5, agg=sum, pool=min | 0.24 | 0.62 | 0.88 | 0.97 | 0.98 | 0.99 | 0.97 | 0.98 | 0.99 | 0.96 | 0.86 |
| 23 \| defl=0.5, agg=sum, pool=max | 0.22 | 0.61 | 0.88 | 0.98 | 0.98 | 1.00 | 0.97 | 0.98 | 0.99 | 0.96 | 0.86 |
| 24 \| defl=0.5, agg=sum, pool=mean | 0.26 | 0.61 | 0.88 | 0.97 | 0.98 | 0.99 | 0.98 | 0.98 | 0.99 | 0.97 | 0.86 |
| 25 \| defl=0.5, agg=product, pool=min | 0.27 | 0.60 | 0.88 | 0.97 | 0.98 | 0.99 | 0.97 | 0.98 | 0.98 | 0.97 | 0.86 |
| 26 \| defl=0.5, agg=product, pool=max | 0.26 | 0.61 | 0.87 | 0.97 | 0.98 | 1.00 | 0.98 | 0.98 | 0.99 | 0.97 | 0.86 |
| 27 \| defl=0.5, agg=product, pool=mean | 0.28 | 0.64 | 0.89 | 0.98 | 0.98 | 1.00 | 0.98 | 0.98 | 0.99 | 0.97 | 0.87 |
| 28 \| defl=0.75, agg=–, pool=– (deflated count only) | 0.21 | 0.56 | 0.88 | 0.98 | 0.98 | 0.99 | 0.97 | 0.98 | 0.99 | 0.96 | 0.85 |
| 29 \| defl=0.75, agg=sum, pool=min | 0.22 | 0.61 | 0.89 | 0.97 | 0.99 | 1.00 | 0.97 | 0.98 | 0.99 | 0.96 | 0.86 |
| 30 \| defl=0.75, agg=sum, pool=max | 0.23 | 0.61 | 0.87 | 0.97 | 0.98 | 1.00 | 0.97 | 0.98 | 0.99 | 0.96 | 0.86 |
| 31 \| defl=0.75, agg=sum, pool=mean | 0.25 | 0.60 | 0.88 | 0.98 | 0.99 | 1.00 | 0.97 | 0.97 | 0.98 | 0.96 | 0.86 |
| 32 \| defl=0.75, agg=product, pool=min | 0.24 | 0.64 | 0.88 | 0.96 | 0.99 | 0.99 | 0.97 | 0.98 | 0.99 | 0.97 | 0.86 |
| 33 \| defl=0.75, agg=product, pool=max | 0.26 | 0.62 | 0.86 | 0.97 | 0.98 | 1.00 | 0.97 | 0.98 | 0.99 | 0.97 | 0.86 |
| 34 \| defl=0.75, agg=product, pool=mean | 0.26 | 0.63 | 0.89 | 0.98 | 0.98 | 0.99 | 0.98 | 0.97 | 0.99 | 0.97 | 0.86 |

Decile of training-set AE count (D1 = 0–5 AEs, D10 = 163–256 AEs)

Mean CV TPR (colour scale 0.0–1.0)

**Table S2:** Top 3 INF configurations per decile (by mean cross-validation TPR); best for each decile in bold.

| **Decile** | **Rank** | **Config idx** | deflator_exp | INF aggregator | pooling | **CV TPR** | **± 95% CI** |
|---|---|---|---|---|---|---|---|
| **1** | **1** | **12** | **0** | **product** | **max** | **0.42** | **± 0.11** |
| 1 | 2 | 13 | 0 | product | mean | 0.42 | ± 0.17 |
| 1 | 3 | 9 | 0 | sum | max | 0.40 | ± 0.12 |
| **2** | **1** | **7** | **0** | **—** | **—** | **0.66** | **± 0.15** |
| 2 | 2 | 27 | 0.5 | product | mean | 0.64 | ± 0.09 |
| 2 | 3 | 15 | 0.3 | sum | min | 0.64 | ± 0.07 |
| **3** | **1** | **15** | **0.3** | **sum** | **min** | **0.91** | **± 0.06** |
| 3 | 2 | 20 | 0.3 | product | mean | 0.89 | ± 0.07 |
| 3 | 3 | 27 | 0.5 | product | mean | 0.89 | ± 0.08 |
| **4** | **1** | **17** | **0.3** | **sum** | **mean** | **0.98** | **± 0.03** |
| 4 | 2 | 31 | 0.75 | sum | mean | 0.98 | ± 0.03 |
| 4 | 3 | 28 | 0.75 | — | — | 0.98 | ± 0.03 |
| **5** | **1** | **17** | **0.3** | **sum** | **mean** | **0.99** | **± 0.01** |
| 5 | 2 | 32 | 0.75 | product | min | 0.99 | ± 0.01 |
| 5 | 3 | 3 | — | sum | mean | 0.99 | ± 0.01 |
| **6** | **1** | **31** | **0.75** | **sum** | **mean** | **1.00** | **± 0.00** |
| 6 | 2 | 30 | 0.75 | sum | max | 1.00 | ± 0.00 |
| 6 | 3 | 29 | 0.75 | sum | min | 1.00 | ± 0.00 |
| **7** | **1** | **17** | **0.3** | **sum** | **mean** | **0.98** | **± 0.01** |
| 7 | 2 | 27 | 0.5 | product | mean | 0.98 | ± 0.01 |
| 7 | 3 | 26 | 0.5 | product | max | 0.98 | ± 0.01 |
| **8** | **1** | **0** | **—** | **—** | **—** | **0.98** | **± 0.04** |
| 8 | 2 | 23 | 0.5 | sum | max | 0.98 | ± 0.04 |
| 8 | 3 | 22 | 0.5 | sum | min | 0.98 | ± 0.04 |
| **9** | **1** | **34** | **0.75** | **product** | **mean** | **0.99** | **± 0.01** |
| 9 | 2 | 30 | 0.75 | sum | max | 0.99 | ± 0.01 |
| 9 | 3 | 5 | — | product | max | 0.99 | ± 0.01 |
| **10** | **1** | **25** | **0.5** | **product** | **min** | **0.97** | **± 0.02** |
| 10 | 2 | 20 | 0.3 | product | mean | 0.97 | ± 0.02 |
| 10 | 3 | 17 | 0.3 | sum | mean | 0.97 | ± 0.03 |

**Figure S3:** Per-decile cross-validation TPR: decile-best INF ensemble components vs best single global INF configuration vs raw-count baseline.

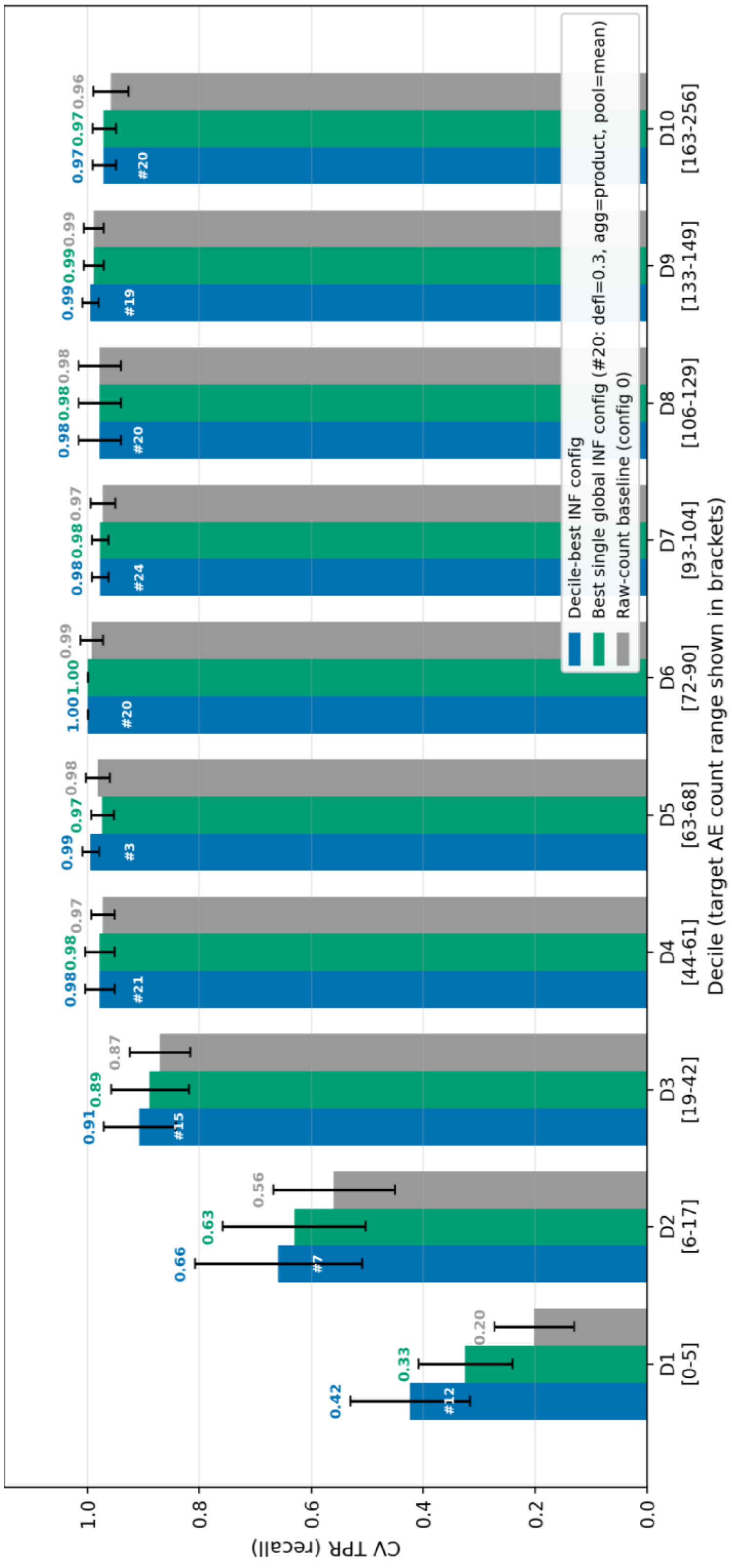

**Figure S4:** Holdout and external test performance across all targets. Left: filtered Hits@10; right: TPR (threshold 0.5), for the INF ensemble, DWPC and original (raw-count) features on the holdout test set (n = 998; solid bars) and the external Soldatos et al. (2018) test set (n = 7,804; hatched bars). Error bars are 95% bootstrap confidence intervals (1,000 resamples).

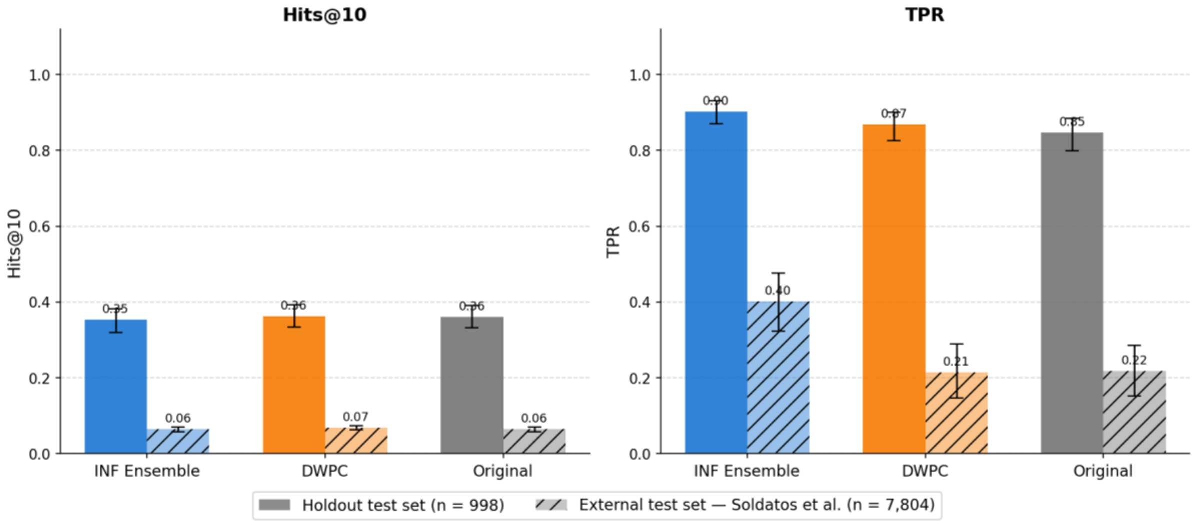


**Table S5:** Random Forest hyperparameters – list of default scikit-learn settings used.

| **Hyperparameter** | **Default Value** | **Brief Description** |
|---|---|---|
| criterion | 'gini' | The function used to measure the quality of a split (e.g., Gini impurity or entropy). |
| max_depth | None | The maximum depth of each decision tree. None means nodes expand until all leaves are pure. |
| min_samples_split | 2 | The minimum number of samples required to split an internal node. |
| min_samples_leaf | 1 | The minimum number of samples required to be at a leaf node. |
| min_weight_fraction_leaf | 0.0 | The minimum fraction of the total sample weight required at a leaf node. |
| max_features | 'sqrt' | The number of features to consider when looking for the best split. 'sqrt' uses √n_features. |
| max_leaf_nodes | None | The maximum number of leaf nodes. None means there is no limit. |
| min_impurity_decrease | 0.0 | A node will be split if this split induces a decrease of the impurity greater than or equal to this value. |
| bootstrap | True | Whether bootstrap samples (sampling with replacement) are used when building trees. |
| oob_score | False | Whether to use out-of-bag samples to estimate the model's accuracy on unseen data. |
| n_jobs | None | The number of CPU cores to use for processing. None means 1; -1 means use all available cores. |
| random_state | None | A seed for the random number generator to ensure reproducible results. |

| | | |
|---|---|---|
| verbose | 0 | Controls the amount of text output during the fitting process. 0 is silent. |
| warm_start | False | If True, reuses the solution of the previous fit and adds more estimators to the ensemble. |
| class_weight | None | Weights for each class, useful for imbalanced datasets. None gives all classes a weight of 1. |
| ccp_alpha | 0.0 | Complexity parameter for Minimal Cost-Complexity Pruning. Higher values increase pruning. |
| max_samples | None | The number of samples to draw from the data to train each base estimator if bootstrap is True. |
| monotonic_cst | None | Enforces a monotonic relationship between specific features and the target. |

**Table S6:** Optimal hyperparameters from cross-validation grid search.

| Feature set | Hyperparameter | | | |
|---|---|---|---|---|
| | max_depth | max_features | min_samples_leaf | min_samples_split |
| Original | None | sqrt | 2 | 2 |
| DWPC | None | sqrt | 1 | 5 |
| INF best for 1st decile | 20 | sqrt | 2 | 2 |
| INF best for 2nd decile | 20 | sqrt | 1 | 2 |
